# Motor history influences behavioral strategy more than response outcome in *Drosophila melanogaster*

**DOI:** 10.1101/2025.07.11.664321

**Authors:** Matteo Bruzzone, Giulio Maria Menti, Marco Dal Maschio, Aram Megighian

## Abstract

In an ever-changing environment, animals continuously implement decision-making processes. This mechanism primarily relies on gathering sensory information through evidence accumulation, which enables the optimization of strategies in uncertain conditions. According to this scheme, the higher the evidence level, the greater the level of accuracy. However, it is not yet clear which, apart from direct sensory inputs, are the other factors that potentially impact strategy selection and their corresponding temporal operating windows in invertebrates. Moreover, it is not guaranteed that maximal accuracy could be the only parameter dictating the decision-making process. To address these questions, we employed a visual stimulation paradigm based on the random dot motion kinetogram (RDMk) with varying coherence levels, applied to tethered *Drosophila melanogaster*. In this scenario, flies tend to turn their heads according to the optic flow’s strength and direction. We observed stereotyped head saccadic-like movements across all the RDMk coherence levels, with greater coherence leading to a higher total saccade number and an increased proportion of correct turns. Importantly, logistic regression analysis revealed that, besides sensory inputs, the behavioral outcome and, consequently, the maximal accuracy, are influenced by other factors, in particular the past motor activity. In conclusion, our data support a scenario where accuracy and reaction time are balanced to reach an optimal response condition.

## INTRODUCTION

In adaptive behavior, animals constantly face scenarios where their actions are critical to their evolutionary success. Some situations involve automatic reflexive responses where relevant sensory inputs trigger a stereotyped motor response with a short reaction time. Other scenarios are more uncertain: in such situations, animals make decisions based on noisy and potentially ambiguous sensory inputs. In these cases, perceptual decision-making can lead to different outcomes, reflecting a tradeoff between computational load, energy costs, and actual benefits. The decision between different choices involves balancing between a slower response with maximal accuracy based on evaluating sensory evidence over time and a faster one with lower accuracy, which mainly relies on *a priori* information. Carefully balancing between these responses is essential, as uncertainty can negatively impact the animals’ fitness by increasing the risk of incorrect responses (errors). For example, it could make animals more vulnerable to predators or reduce their ability to locate prey [Salles *et al*., 2020]. In such cases, the animal’s perceptual decision-making process may benefit from accumulating and integrating sensory information over time, which can act as a bias to improve accuracy. However, even under conditions that should facilitate accurate decisions, individuals make errors. This discrepancy can be attributed to the fact that achieving optimal accuracy often requires time. In natural settings, however, rapid responses are typically favored over perfectly accurate ones, as they can yield satisfactory outcomes with lower energetic costs. Consequently, a trade-off arises between achieving optimal accuracy and response promptness [Bogacz, 2007; Gold and Shadlen, 2007].

Strategies that reduce the computational load (i.e., processing the minimal amount of information) and shorten decision-making time while still leading to adaptive behavior are termed heuristics [Gigerenzer and Gaissmaier, 2011]. Heuristics provide quicker, efficient solutions that, although not achieving the theoretical maximal accuracy [Dale, 2015], are practical and well-suited to the challenges of real-world settings, and thus may match the optimality for a given scenario. This approach may draw on the combination of sensory perception with the recent history of observed stimuli [Akrami *et al*., 2018; Fisher and Whitney, 2014], experienced rewards [Abrahamyan *et al*., 2016; Gupta *et al*., 2024; Urai *et al*., 2019], and current motor history [Hwang *et al*., 2017; Dragomir *et al*., 2020]. These strategies have been observed across a range of animal species, mainly vertebrates [Bahl and Engert, 2020; Hanks and Summerfield, 2017], but little is known about whether similar strategies are also employed by invertebrates. Studying whether heuristics are present in invertebrates and how they operate could unravel their significance throughout evolution. Moreover, the simplicity of invertebrates’ nervous systems could help in investigating the neurobiological basis of these cognitive mechanisms. The most studied invertebrate species is *Drosophila melanogaster* (*Dmel*), and previous research has shown that *Dmel* can accumulate evidence during olfactory decision-making [DasGupta *et al*., 2014; Groschner *et al*., 2018] and use heuristics during spatial learning [Meda *et al*., 2022]. However, it is unclear whether *Dmel* is capable of integrating sensory perception with its motor history and if this can play an active role in the perceptual decision-making process, particularly when the environment is characterized by uncertainty. To address this question, we devised an experimental paradigm where the level of uncertainty can be effectively tuned on the features of sensory stimuli. We took advantage of “Random Dot Motion” kinetogram (RDMk), a consolidated visuomotor stimulation paradigm for investigating evidence accumulation, where the relative contribution of noise and uncertainty is modulated by varying the fraction of dots moving coherently along a set direction with respect to the component of dots moving incoherently in random directions (Gold & Shadlen, 2007). This type of stimulation drives a sensorimotor integration task where motion direction discrimination prompts a motor response according to the direction of global motion (Gold & Shadlen, 2007). By selecting the most appropriate motor output reflecting the coherent direction of the dots, it is possible to study an organism’s motion discrimination. In our experiment, we focused on *Dmel*’s head movements as a proxy of visually evoked motor response [Fox and Frye, 2014; Keleş *et al*., 2019].

We found that flies are capable of integrating visual evidence, thereby modulating their accuracy, which is defined as the rate of correct motor responses. We noted that this gathering of sensory information before a given movement is influenced by the motor history spanning more than the previous movement. Interestingly, our analysis shows that accuracy is significantly enhanced by integrating sensory information with the history of motor outcomes and the information about their correctness.

## RESULTS

### Flies perform stereotypical nystagmus-like head movements in response to RDMk stimulation

To investigate the motor responses during an evidence accumulation task, we employed an RDMk paradigm. To track the behavior during visual stimulation, we devised an apparatus where individual flies are tethered to a pin while facing a panoramic screen displaying the stimuli. Two infrared LEDs illuminated the fly, and an infrared camera, positioned below the fly, recorded its head movements (Fig. 1A - left). In response to RDMk stimulation, flies exhibited stereotypical head movements, we previously named these head optokinetic nystagmus (HOKN) [Menti *et al*., 2024], characterized by two alternate components: a slow tracking phase according to the perceived dots’ motion direction which is followed by a fast phase (a “saccade”, see also [Cellini and Mongeau, 2021]) in the opposite direction (Fig. 1A – right). During the 20 minutes of stimulation, flies performed 838.95 ± 151.57 head movements (mean ± standard deviation, n:21). In response to the stimulation, flies exhibited head angular turns of approximately 6 degrees of amplitude during the slow phase, with the head turning direction peaking according to the RDMk coherent component moving in a set direction (Fig. 1B); meanwhile, the fast phase was characterized by a peak angular velocity of approximately 118 deg/s (Fig. 1C) in the opposite direction of the movement performed during the slow phase. While we did not observe any significant modulation on these two HOKN kinematic parameters depending on the stimulus coherence, the number of head movements showed a significant increase with the level of coherence (Fig. 1D), suggesting that flies are able to perceive differences in the motion coherence level of the stimulus.

**Figure 1.**
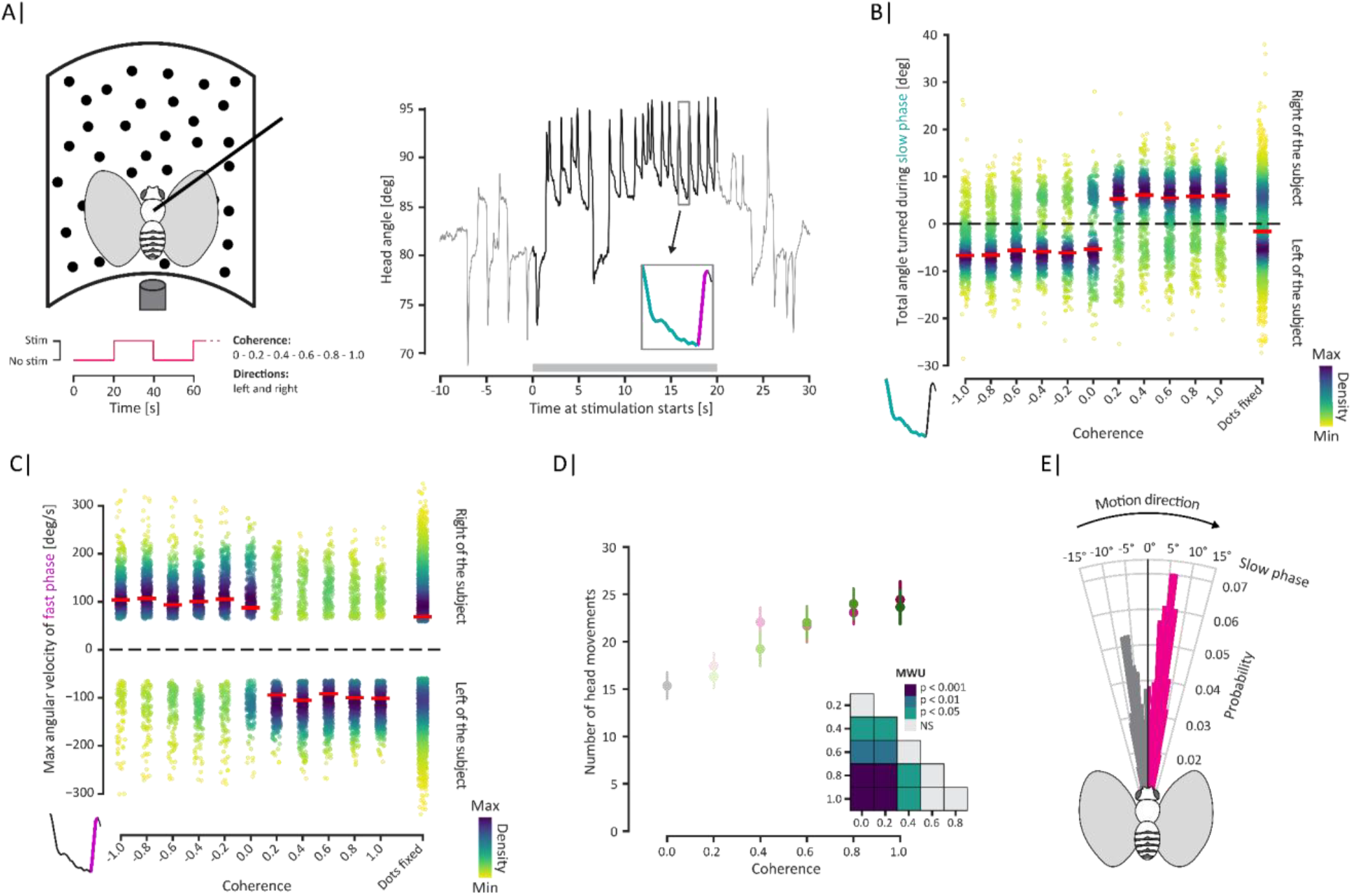
**A)** Left: The fly is suspended above an infrared camera, while facing a screen on which visual stimuli are presented. Stimulations (STIM) consist of 20-second phases with a fraction of the presented dots moving in one of the two directions - either toward the left or the right of the still Dmel midline, while the remaining dots move randomly. Stimulations were interleaved by 20-second phases with all the dots kept still (NO STIM). Right: Typical behavioral recording. The magenta box highlights a head movement, showing slow (stimulus tracking) and fast phases. **B)** Total angle turned during the slow phase (teal colored trace in the bottom left corner) as a function of the coherence level. Red lines are medians. **C)** Maximum angular velocity during the fast phase (magenta colored trace in the bottom left corner). Red lines are the median. **D)** Average number of head movements per subject. For coherence levels greater than 0, data for both left and right directions (green and magenta, respectively) are shown. Mann-Whitney U (MWU) test results comparing coherence levels are reported in the half-matrix. All error bars indicate the standard error of the mean (s.e.m.), n = 21. **E)** Probability distribution of the angle turned during the slow phase pulled across the coherence levels. In magenta, the slow phases aligned with the motion direction of the dots.

### Accuracy increases across time and movements

Looking at the distributions of the behavior responses, we noticed that while most stimulations led to movements co-directional with the dots’ motion direction during the slow phase (Fig. 1E - magenta), a non-marginal number of outcomes were instead motions in the opposite direction (i.e., slow-phase movements opposite to the dots’ motion direction). The number of these *non-directional* head movements showed a decreasing trend as the coherence level rose, but never completely resolved, even at maximal coherence level, so constraining the accuracy (Fig. 1E – gray and Fig. 2A). This suggested that other factors could participate and influence the decision on the motor outcome, and eventually impact the observed accuracy. The dependence of the response on the persistence of the stimulus presentation is considered a key aspect during the integration of evidence, as well as the refinement of this same response during consecutive accumulations. Therefore, we explored whether flies could tune their accuracy throughout the stimulation, similar to what is observed in other species [Dragomir *et al*., 2020; Tanimoto and Kimura, 2019]. We defined “correct movements” as those where the slow phase was coherent with the direction of the dots’ movement, and a*ccuracy* as the proportion of correct movements to the total number of head movements (co-directional + non-directional). In our configuration, the RDMk was continuously presented for 20 seconds with a constant coherence level: the cumulative turn angle increased with the stimulus coherence, supporting the conclusion that flies respond to RDMk as motion of varying strengths (Fig. 2B). Evaluating this aspect, we found that flies indeed increase their accuracy over time. Such trend was particularly pronounced at coherence levels 0.8 and 1.0, where accuracy plateaued at approximately 80% within a few seconds (Fig. 2C). Accuracy also increased with the number of performed movements (Fig. 2D) and persisted after the stimulation ceased, displaying a gradual decline over the successive movements (Fig. 2E). Interestingly, we noticed that the first movement after the stimulus onset always resulted to be random, irrespective of stimulus intensity, and its accuracy was independent from the time at which it was performed (Fig. S1).

**Figure 2.**
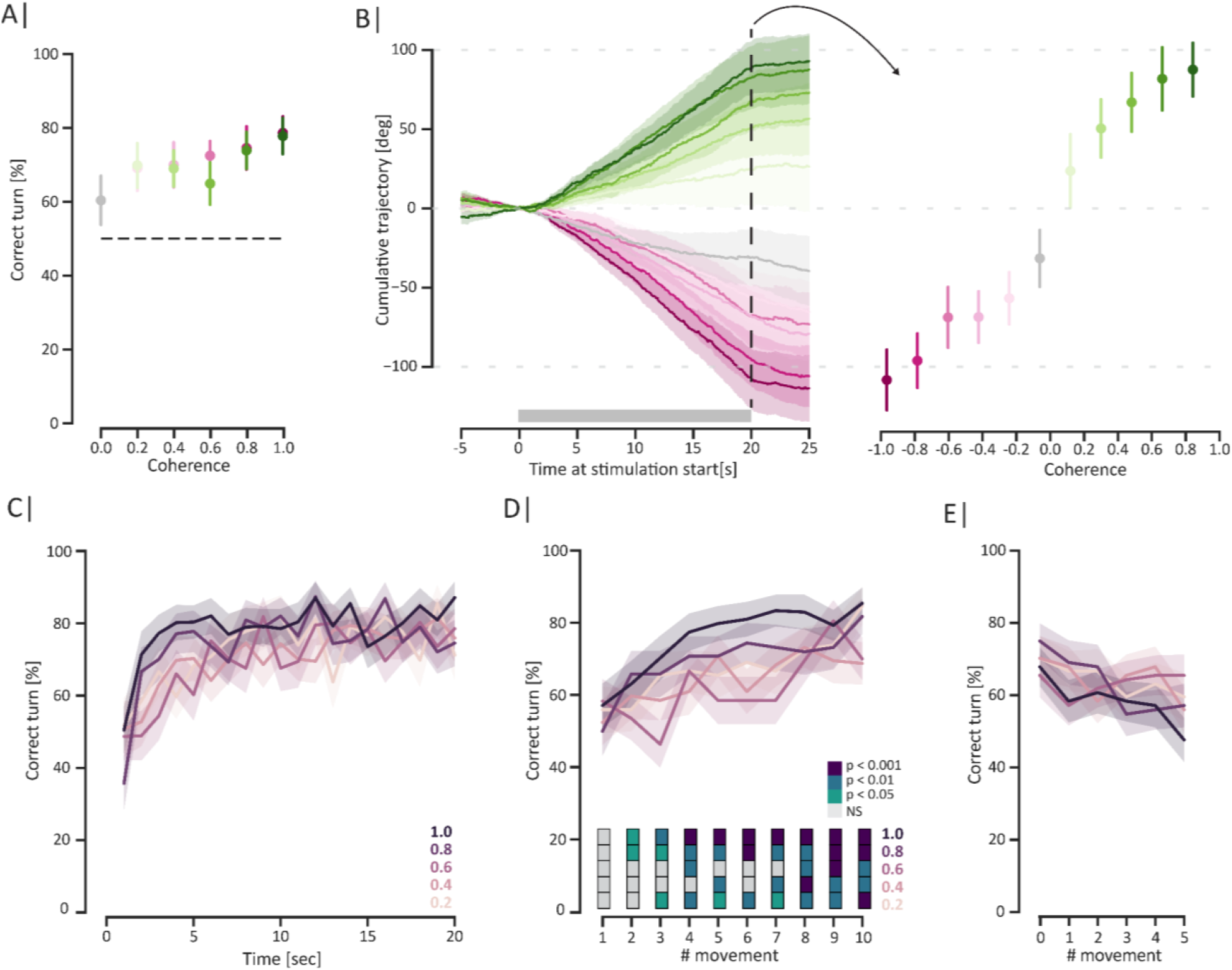
**A)** Proportion of correct turns as a function of coherence level, averaged across subjects. For coherence levels above 0, data for both left and right directions (green and magenta, respectively) are shown. The dashed line indicates the chance level. **B)** Left: Average cumulative angle turned as a function of coherence level during the 20s of stimulation (grey bar). Right: Average total angle turned at the end of the stimulation. **C)** Proportion of correct turns as a function of time for different coherence levels. **D)** Fraction of correct turns as a function of head movements from the start of the stimulation at different coherence levels. (Wilcoxon signed-rank test). **E)** Fraction of correct turns as a function of head movements from the end of the stimulation at different coherence levels. All error bars and shaded intervals indicate the s.e.m. n: 21.

These results agree with the presence of an accumulation process that spans over a timescale longer than single movements.

### Sensory input and previous motor choices are integrated into a decision

The performance of actual observers has been shown to deviate from the ideal one expected from maximal accuracy [Gardner, 2019]. This is due to various factors, such as past rewards and previous motor choices [Abrahamyan *et al*., 2016; Gupta *et al*., 2024; Urai *et al*., 2019]. These dependencies on prior history are especially evident when the stimulus is presented randomly and with varying intensities [Urai *et al*., 2019]. In our case, the accuracy measured experimentally never exceeded the 80% level (Fig. 2A). We then asked whether, and to what extent this process could be influenced by motor history, considering in particular two aspects: *i*) the previous motor action and *ii*) its correctness to the visual stimulation features. We found that both the previous motor choice (i.e., left or right) and the previous outcome (i.e., wrong or correct) significantly impact the accuracy (Fig. 3A). Specifically, considering appropriate stimulus-behavior sequences, we found that the accuracy increases significantly when the current motor action was in the same direction as the previous one (Fig. 3A - Choice). Moreover, data showed higher accuracy when the previous choice led to a successful outcome, i.e., the outcome from the previous stimulation was in the same direction as the dots’ motion (Fig. 3A - Outcome).

**Figure 3.**
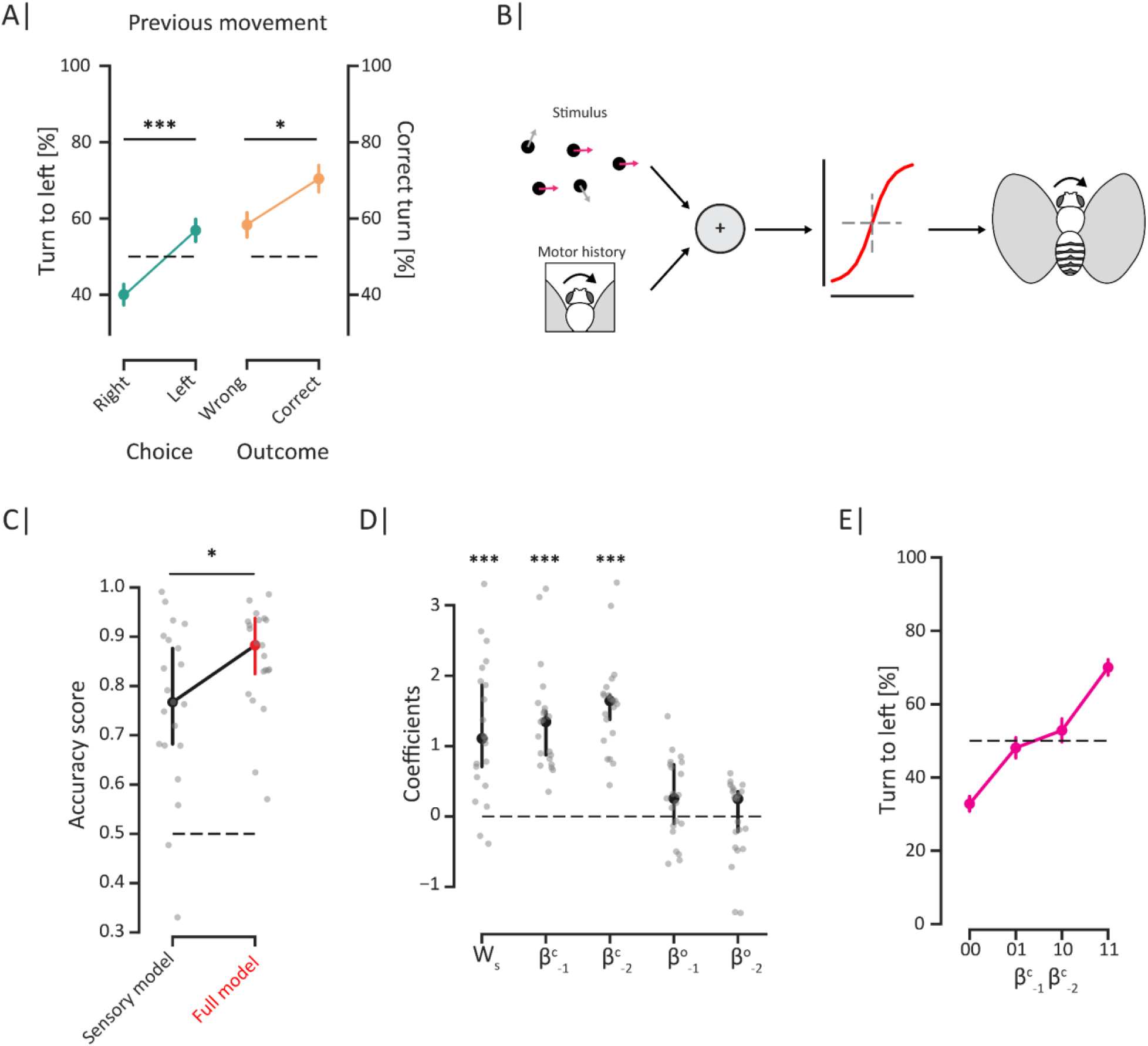
**A)** Proportion of turns to the left as a function of the previous movement direction (left or right) or outcome (correct or wrong relative to the dots’ motion direction), averaged across all subjects and coherence levels. The dashed line indicates the chance level. Asterisks denote statistical differences (Mann-Whitney U test - two-sided, *: p<0.05, ***: p<0.001). **B)** Cartoon of the proposed logistic model. **C)** Accuracy scores of the logistic classifier using the sensory model versus the full model, averaged across subjects. The dashed line indicates the chance level. The asterisk denotes a statistical difference (Mann-Whitney U test - one side, *: p<0.05). **D)** Average values of the coefficients in the full model, averaged across subjects. WS denotes the visual stimulus contribution, β^c^ denotes the motion choice contribution, and β° denotes the turn outcome contribution. Asterisks denote significant deviations from 0 (Wilcoxon signed-rank test, ***: p<0.001). **E)** Proportion of turns to the left as a function of the two previous movement choices, averaged across subjects. 0 represents a right turn and 1 represents a left turn. All error bars and shaded intervals indicate the s.e.m. except in C) and D), which denote a 95% confidence interval.

To model this aspect, we developed a multivariate logistic classification model taking into account both the stimulus and the motor history (Fig. 3B). The model incorporated, along with the sensory evidence contribution, *W*_*S*_, both a motor choice component represented by a coefficient β^*c*^ (left or right), and a second component to account for the motor outcome represented by the coefficient β^*o*^ (correct or incorrect). Anticipating that such contributions could operate in different temporal scales, as reported in other species [Dragomir *et al*., 2020], we analyzed the impact of up to five previous motor events 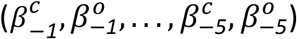. Our model shows that the two previous movements significantly impact accuracy (Fig. S2A). The model, including the sensory input and the two previous movements (parametrized by 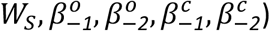), was better at capturing the subjects performance, achieving a median accuracy score of 0.888 (95% CI [0.812, 0.939]), compared to the simpler model, comprising only the sensory information, which had a median accuracy score of 0.767 (95% CI [0.682, 0.893]) (Fig. 3C, Fig. S2B). Remarkably, by looking at the corresponding weights, it emerges that the previous choices β^c^ (left or right) have a much stronger predictive power than previous motor outcomes β° (wrong or correct), highlighting a strong bias on the behavioral choice exerted by the motor history rather than by the correctness (Fig. 3D). In our view, this motor bias could represent an *internal* bias of the subject, which acts irrespective of the present/current visual stimulation, and influences the accuracy of the subject biasing toward the repetition of the previous choices (Fig. 3E). Interestingly, the bias got stronger as time proceeded (Fig. S2C), as expected in an information accumulation process.

## DISCUSSION

In complex and mutable environments, animals constantly face scenarios where decisions determine their survival. This relies primarily on sensory input elaboration through a cognitive process known as perceptual decision-making, which transforms potentially noisy sensory information into behavioral outcomes (i.e., motor outputs). This process has been studied in more detail in humans, primates, and rodents [Hanks and Summerfield, 2017]. However, studies conducted on animals that are more distant from humans have highlighted how this could be a shared mechanism across different sensory modalities [Najafi and Churchland, 2018]. The relative maintenance of this process in both simple and complex animals reveals its fundamental role in adaptive behavior.

Dealing with uncertain sensory inputs, it is speculated that perceptual decision-making operates by balancing the accuracy of the response and the reaction time [Gardner, 2019]. Both aspects play a fundamental role in adaptive response and the evolution of behavior, intended as “the internally coordinated responses (actions or inactions) of whole living organisms (individuals or groups) to internal and/or external stimuli, excluding responses more easily understood as developmental changes” [Levitis *et al*., 2009]. Consequently, a trade-off arises between achieving optimal accuracy and timely responses [Bogacz, 2007; Gold and Shadlen, 2007]. Finding this context-dependent optimal balance would correspond for any organism in a given situation to reaching response optimality, not necessarily implying absolute accuracy. Moreover, in this context, the term “error” becomes nonsensical unless used for defining the distance between the real observed maximal accuracy and the theoretical one (which is more or less close to 100%). To put it bluntly, this equals saying that an organism is bound to the optimality of its responses, independently of the error. However, optimality can still be quantified, identifying upper and lower limits in the range of the enacted responses - like the 0-80% interval in our data - which will depend on the pressures (internal or external) threatening or actively shifting the system’s equilibrium (e.g., the fly trying to stabilize its course and adjusting its flight) by a measurable amount.

Another key concept of this process is evidence accumulation, which involves consolidating sensory information over time before a decision is made. While perceptual decision-making has been studied across various animal models reviewed in [Hanks and Summerfield, 2017] and sensory modalities, research in *Drosophila* has mostly focused on odor-based tasks [DasGupta *et al*., 2014; Groschner *et al*., 2018].

In this study, we used the established RDMk paradigm, where visual sensory “noise” can be tuned, to explore how flies engage in visual evidence accumulation. Our findings reveal that, akin to more complex organisms, flies can accumulate visual evidence. More coherent sensory inputs result in more accurate motor outputs, suggesting that the relative magnitude of the sensory input signal-to-noise ratio influences behavioral choices, with stronger accumulated evidence leading to less erratic outcomes - a result consistent with previous findings that stimulus strength impacts both decision accuracy and response time [Gold and Shadlen, 2007]. Indeed, in our case, stimulus strength influences not only the accuracy but also the number of responses, while the relationship between the response time and accuracy does not seem to be affected. Here, we see that flies integrate sensory evidence over time, with higher coherence yielding greater accuracy within the first seconds, while weaker stimuli lead to a smoother increase in accuracy. However, despite initial differences at the beginning of the stimulation, flies never achieved optimal accuracy, and by the end of the stimulation, accuracy became consistent across varying stimulus intensities. This suggests two key points. First, the stimulus signal-to-noise ratio is not the sole factor driving the observed behavior, although it plays a significant role, especially at the onset of the stimulation [Gardner, 2019; Palmer *et al*., 2005]. Second, even in optimal conditions, with minimal or no noise, flies exhibit some deviations, suggesting the presence of additional factors that drive the behavior and, consequently, dampen the level of accuracy.

The dynamics of decision-making processes have been extensively studied, with models like the bounded leaky integrator and drift-diffusion being widely used to characterize evidence accumulation in these processes [Hanks and Summerfield, 2017]. While these models can achieve near-optimal performance under certain conditions, real-world decision-making often deviates due to the complexity and noise in the environment. Again, this outcome is adaptive and results from the tradeoff between response time and accuracy [Bogacz, 2007; Gold and Shadlen, 2007]. Like other animals, flies appear to adopt strategies that prioritize faster decision-making by focusing on less information, even at the cost of reduced accuracy. Such heuristics have been observed in mammals during evidence accumulation tasks, including dependencies on stimulus history, reward history, or motor history. In the context of the RDMk paradigm, where binary outcomes are influenced by varying sensory weights (coherence), logistic classification provides a robust analytical framework, and it has been successfully applied across a range of animal studies to model decision-making behavior [Gardner, 2019]. In our study, logistic classification proved effective in describing the behavioral responses of flies. We found a significant influence of motor history on responses, suggesting that flies employ strategies extending beyond immediate motor outputs. In this scenario, flies employ a behavioral strategy that does not lead to the maximization of accuracy but rather is biased by the execution of a consolidated motor program, at least to a certain degree.

Why flies utilize this strategy is a matter of speculation. Certainly, the response to this question may lie in the relative complexity of flies’ nervous system with respect to higher mammals, which makes it more difficult to reach optimality through a time-consuming process (and possibly a processing activity which may be too heavy for the capacity of this nervous system). In this view, heuristics could represent a basal yet sophisticated layer of cognitive decision-making processes in the organisms’ nervous systems throughout evolution.

## METHODS

### Fly strains

We used Berlin-K female flies aged 3-5 days old. Flies were reared at a controlled temperature of 25°C under a 12-hour light/12-hour dark cycle with ad libitum access to food and water. Experiments were performed within 6 hours of light onset.

### Tethered paradigm

Subject preparation was carried out as described previously (Menti *et al*., 2024). Briefly, subjects were cold anesthetized in ice and placed individually on a Peltier mounting stage maintained at 4°C. A stainless steel needle (BSTEAN, Shenzhen Hemasi E-Commerce Co., Ltd., PRC) was attached to the fly using UV-curable resin (DecorRom, Shenzhenshi Baishifuyou Trading Co., Ltd., PRC – Model Number: ASKU0035). The pin was placed on the upper part of the thorax at an angle of approximately 60°, projecting forward. Flies were allowed to rest for at least 15 minutes before the experiments.

Following acclimatization, subjects were placed in a custom-built arena with black walls to block ambient light during the experiment. The subject, secured to the pin, was mounted on a vertical support and suspended above an Infrared Camera (MQ003MG-CM, Ximea GmbH, Germany) recording at 200 fps. Two IR (850 nm) LEDs (M850L3, Thorlabs Inc., US) illuminated the subject from the sides. A custom-made bent screen of parchment paper was placed in front of the subject. The screen covered ±90° in azimuth and ±45 degrees in altitude. A projector (Lightcrafter 4500, Texas Instruments Inc., US) was placed behind the parchment screen and was used to display visual stimuli at 60 Hz. The video recording and the stimulation were synchronized using a custom script based on Stytra (Štih et al., 2019), running on an HP precision 3660; 12th Gen Intel(R) Core (TM) i7-12700 2.10 GHz and 32 GB RAM.

### Visual stimulation

Random dots of 1 mm size were presented for 20 seconds, followed by 20 seconds of pause (i.e., dots fixed). Six (6) levels of coherence were presented: 0.0, 0.2, 0.4, 0.6, 0.8, 1.0 in two directions, respectively, the subjects’ left and right. The lifespan of the dots was set to a maximum of 0.3 seconds. Each combination of coherence and direction was presented two times. The total protocol lasted 20 minutes.

### Tracking and head movements identification

Tracking was conducted offline using the standalone MATLAB program ‘Flyalyzer’ [Rauscher and Fox, 2021]. Subjects with poor illumination and those moving less than 90% of the total experiment duration were excluded from the analysis. Head time series were initially filtered using a Butterworth low-pass filter with a 5 Hz cutoff frequency, followed by smoothing with a Gaussian filter (σ = 2) to remove tracking artifacts and background noise. Peak detection was performed in two stages using the ‘find_peaks’ function from the ‘scipy’ library [Virtanen *et al*., 2020]. First, a threshold was calculated by taking the median of the absolute angular velocity (i.e., the absolute values of the head movement derivative) and dividing it by 0.6745. This value was derived from a previous work [Mongeau and Frye, 2017] and proved effective with our data. Peaks were considered valid whenever they fell between one and ten standard deviations. Peak prominence, a critical factor in distinguishing head movements and sudden position changes, was used to refine peak detection. We calculate the lower and upper prominence thresholds as the 0.01 and 0.99 quantiles of the previously identified peaks. Peak identification was then repeated, including the prominence thresholds.

### Logistic classification

To determine how the direction of the head turns depends on: *i*) the current stimulus, *ii*) the previous head turn, and *iii*) the outcome of the previous head turn, we designed a logistic multivariate regression model. This model describes the probability of turning in one direction as:

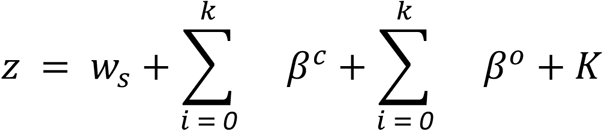

*w*_*s*_ is the weight associated with the stimulus, β^*c*^_*i*_ is the direction of the *i* previous head turn, and β^*o*^_*i*_ is the outcome (i.e., whether it was in the same or opposite direction as the stimulus) of the *i* previous head turn. *K* denotes the stimulus prior bias. The dots had an equal 50% probability to move in either direction, so no stimulus prior bias was present. As a result *K* was equal to 1 and therefore negligible. We fitted the model using the ‘statsmodels’ module [Seabold *et al*., 2010] in Python for each subject and then averaged across them. To determine the optimal number of previous movements to include in the model, we evaluated the cross-entropy loss function (CELF) for partial models with sequentially increasing numbers of previous movements, up to 5 (Fig. S3A). The ideal number of motor covariates was identified as the point with the maximum distance from the y = x line (Fig. S3A) [Satopää *et al*., 2011].

## ACKNOWLEDGMENTS

The authors thank Anita Cattapan and Giuseppe Beltrame, who contributed to the collection of the data presented in this paper.

## Funding

The work was funded by the 2020 PRIN program from the Italian Ministry for the University and Research (MIUR), assigned to project n.° 2020529PCP_001 (“*Free energy principle and the brain: neuronal and phylogenetic mechanisms of Bayesian inference*” - P.I.: Prof. Ettore Ambrosini - University of Padova, Department of Neuroscience). Additional funding came from the DOR research funds assigned to Prof. Marco Dal Maschio and Prof. Aram Megighian by the University of Padova’s Department of Biomedical Sciences.

## Authors’ conflict of interest statement

The authors declare no conflict of interest.

## Data availability

The relevant datasets are available in the Zenodo repository “*Motor history influences behavioral strategy more than response outcome in Drosophila melanogaster” (Bruzzone et al*., *2025) - Dataframes and logs*” at https://doi.org/10.5281/zenodo.15861476. The analysis script can be shared by the authors upon request.

## Authors’ contribution

MB, GMM, and AM set up the experimental apparatus. MB and GMM supervised or carried out the experiments. MB analyzed the data, with feedback from all other authors. MB and GMM drafted the article. All authors reviewed and contributed to the present version of the article.

---

**Figure S1.**
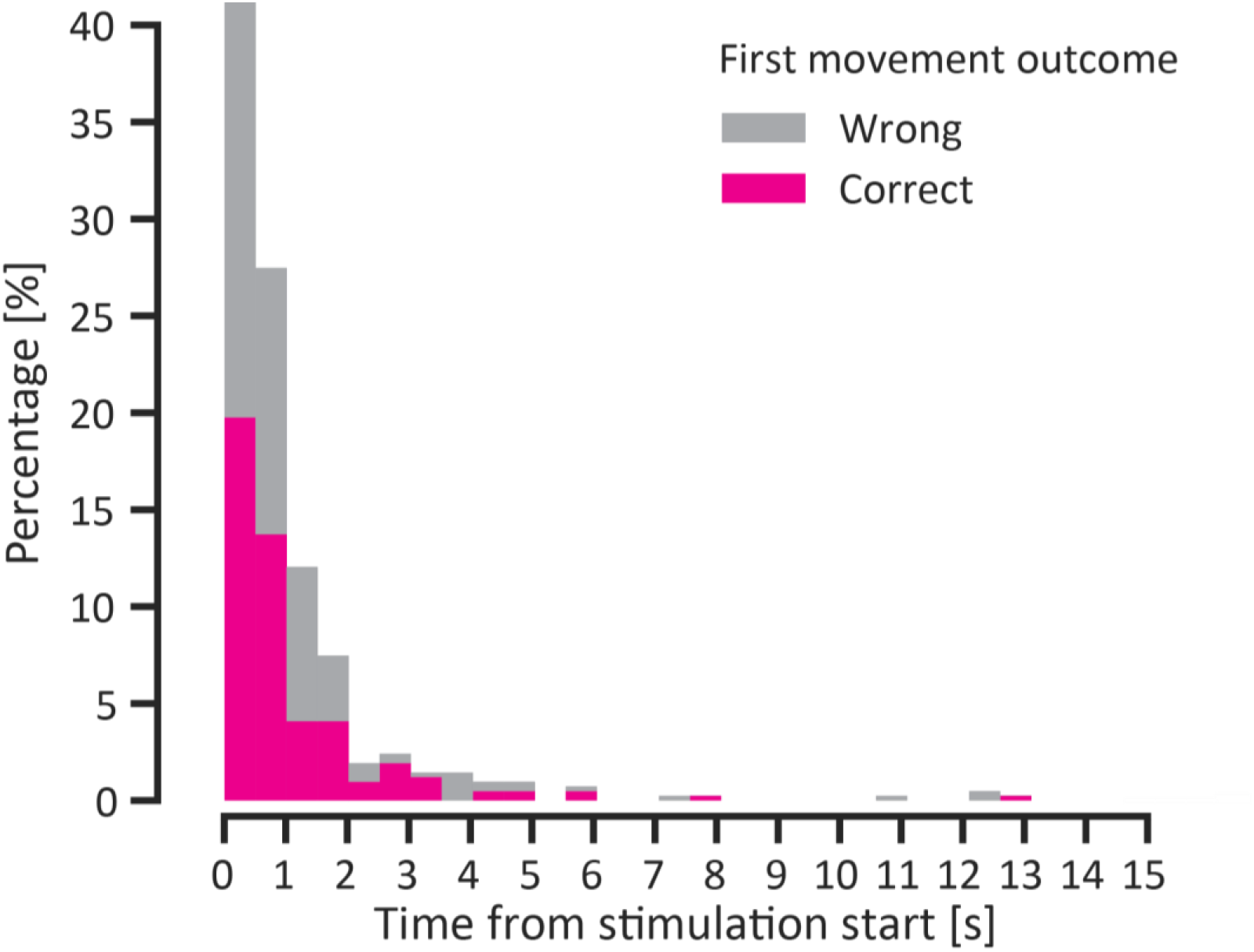
Outcome of the first movement performed from the start of the stimulation.

**Figure S2.**
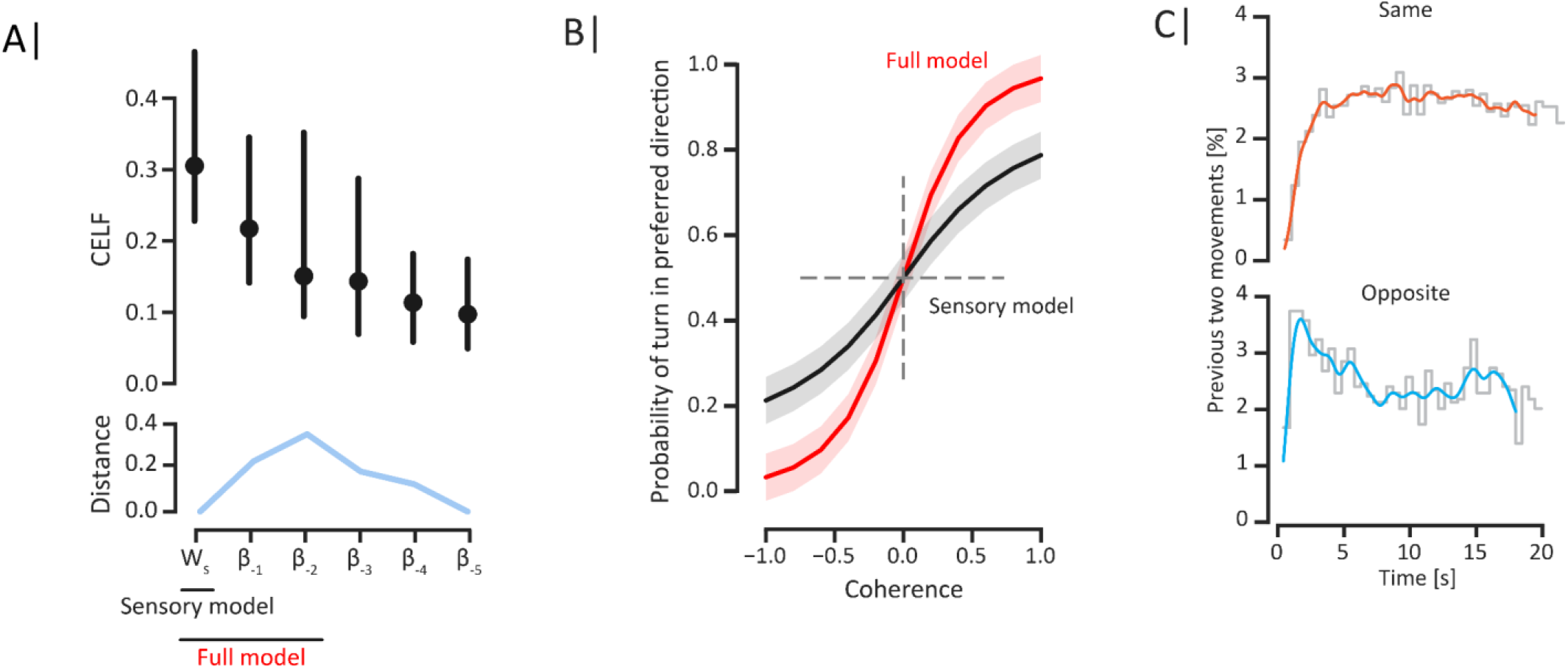
**A)** Cross Entropy Loss Function (CELF) of distinct logistic models with an increasing number of components. The optimal number of components for the full model was determined by identifying the point farthest from the x = y line, representing the ‘knee’ of the CELF curve. W_S_ denotes the visual stimulus contribution, and β denotes the motor contribution, including both choice and outcome. Subscript numbers indicate the previous movement numbers. Error bars indicate a 95% confidence interval. B) Logistic sigmoids of the Sensory model and the Full model. The shaded area indicates s.e.m. C) Frequency of the two consecutive movements being in the same (top) and the opposite direction (bottom) as a function of time.

## Notes

https://doi.org/10.5281/zenodo.15861476

### Competing Interest Statement

The authors have declared no competing interest.

